# Macroscopic to Ultrastructural Analyses Identify the Loss of Myofibrils as the Primary Mediator of Muscle Fiber Atrophy in Aging and Disuse

**DOI:** 10.64898/2026.02.16.706166

**Authors:** Ramy K. A. Sayed, Anthony N. Lange, Hector G. Paez, Jamie E. Hibbert, Marius Meinhold, Corey G.K. Flynn, Matilde B.Z. Vergara, Isabell Dobrzycki, David J. Wrucke, Carlos S. Zepeda, Jessica J. James, Christopher W. Sundberg, Troy A. Hornberger

## Abstract

**Background:** Aging and disuse are two of the most clinically relevant conditions associated with the loss of skeletal muscle mass, yet the ultrastructural adaptations that drive these losses remain poorly defined. In particular, it is unclear whether radial atrophy of muscle fibers is driven by a reduction in the size of the existing myofibrils, and/or the loss of myofibrils. Accordingly, the objective of this study was to define the macro-to-ultrastructural adaptations that mediate aging- and disuse-induced loss of muscle mass.

**Methods:** Skeletal muscle structure was assessed at the macroscopic, microscopic, and ultrastructural levels in humans and mice. In humans, magnetic resonance imaging was used to quantify knee extensor muscle volume and cross-sectional area (CSA) in young (19 - 40 years) and old (65 - 84 years) adults, and vastus lateralis biopsies were analyzed for microscopic and ultrastructural adaptations using immunohistochemistry and fluorescence imaging of myofibrils with image deconvolution (FIM-ID). Parallel studies were performed in young (4 months) and aged (24 months) mice, along with the use of unilateral hindlimb immobilization to model disuse.

**Results:** Aging led to a robust loss of skeletal muscle mass that was mediated by coordinated macro-to-ultrastructural adaptations. In humans, aging reduced knee extensor muscle volume (34%, *P <* 0.005) and CSA (32%, *P* < 0.001) in a sex-independent manner, and these effects were associated with radial atrophy of SERCA1-positive fibers (23%, *P* < 0.05). Ultrastructural analyses revealed that the radial atrophy was driven by a reduction in the number of myofibrils per fiber (23%, *P* < 0.05) without changes in myofibril CSA. In mice, aging produced similar macro-to-ultrastructural adaptations in various flexor muscles; however, radial atrophy of the highly glycolytic/Type IIb fibers, which are not present in human limb muscles, was also associated with a decrease in the CSA of the myofibrils (9%, *P* < 0.005). We also determined that disuse led to radial atrophy of SERCA1-positive fibers (24%, *P* < 0.001), and this was mediated by a decrease in both the number (22%, *P* < 0.005) and size of the myofibrils (4%, *P* < 0.05). Notably, the results also revealed that the magnitude of the disuse-induced adaptations was significantly blunted with aging.

**Conclusion:** This study identifies the loss of myofibrils as a central and conserved mediator of the radial atrophy of muscle fibers that occurs in response to disuse and aging, while also highlighting smaller context-dependent contributions that can arise from changes in myofibril size.

## Introduction

Skeletal muscle is the largest internal organ in the human body, accounting for approximately 35% of the total body mass [1]. Beyond its primary role in locomotion, skeletal muscle performs a variety of other functions, including thermoregulation, support of respiration, maintenance of metabolic homeostasis, and endocrine signaling through the secretion of myokines [2–4]. Given these multifaceted functions, it should not be surprising that the maintenance of skeletal muscle mass has become widely recognized as being critical for overall health and quality of life [2, 5, 6]. Indeed, the loss of skeletal muscle mass has been associated with an increased risk of type II diabetes, cardiovascular diseases, and all-cause mortality [7–9]. Thus, understanding the mechanisms that regulate skeletal muscle mass is crucial, as this knowledge will guide the development of therapies for preventing the loss of muscle mass and improving public health.

At the most fundamental level, changes in skeletal muscle mass arise from alterations in the size and/or number of its structural components. These components are organized in a hierarchical manner that can be examined at three levels: the macroscopic level (visible without magnification), the microscopic level (observable using standard light microscopy), and the ultrastructural level (requiring high-resolution imaging techniques such as electron microscopy or super-resolution light microscopy) [10, 11]. Generally speaking, the macroscopic level encompasses the anatomical properties of the whole muscle, the microscopic level reflects features of the individual muscle fibers, and the ultrastructural level pertains to the properties of the myofibrils that are found within the fibers. Importantly, the hierarchical organization of the structures predicts that adaptations at lower levels propagate upward in a predictable manner. For instance, changes at the level of the myofibrils should lead to predictable changes at the level of the muscle fibers, which, in turn, should lead to predictable changes at the whole-muscle level.

One of the most clinically relevant contexts for studying the mechanisms that regulate muscle mass is aging. Specifically, with advancing age, skeletal muscle undergoes a progressive loss in mass through a process known as sarcopenia [12, 13]. At the macroscopic level, numerous studies have demonstrated that sarcopenia is marked by a decrease in whole muscle cross-sectional area (CSA), and that this effect has been attributed to coordinated changes at the microscopic level, including both the loss of muscle fibers (i.e., hypoplasia) and a reduction in the CSA of the muscle fibers (i.e., radial atrophy) [14, 15].

Disuse represents another clinically relevant context in which skeletal muscle mass is profoundly reduced [16, 17]. Indeed, a variety of models in humans and lower organisms have consistently shown that disuse leads to a decrease in whole muscle CSA [11, 18, 19]. However, unlike with aging, the disuse-induced decline in whole-muscle CSA does not appear to involve hypoplasia. Instead, most studies indicate that the decrease in whole muscle CSA is largely driven by radial atrophy of existing muscle fibers [11, 20, 21].

Although the induction of radial atrophy that occurs with disuse and aging has been recognized since the 1970’s [22, 23], the underlying ultrastructural adaptations that drive this phenomenon remain poorly defined. In fact, even the most basic questions, such as whether the radial atrophy of the muscle fibers is driven by a decrease in the CSA of the myofibrils and/or a reduction in the number of myofibrils, remain largely unresolved. Thus, the primary objective of this study was to identify the macro-to-ultrastructural adaptations that mediate the aging- and disuse-induced loss of muscle mass.

## Methods

### Participants and ethical approval

Ten young males (19-40 yrs), eight young females (21-28 yrs), nine old males (69-84 yrs), and seven old females (65-72 yrs) volunteered and provided their written informed consent to participate in this study. Participants underwent a general health screening that included an assessment of body composition and thigh lean mass with dual X-ray absorptiometry (Lunar iDXA; GE, Madison, WI). Participants were excluded from the study if they were taking medications that may affect muscle mass (e.g., hormone-replacement therapies, glucocorticoids, etc.). All participants were apparently healthy, community-dwelling adults free of any known neurological, musculoskeletal, or cardiovascular diseases. Following the general health screening, participantsreported to the laboratory on two occasions: once for a muscle biopsy of the vastus lateralis, and again to quantify the size of the knee extensor muscles with proton magnetic resonance imaging (MRI). The anthropometrics of the young and old participants are shown in **Table 1**. All experimental procedures were approved by the Marquette University Institutional Review Board (Protocol Number HR-2945) and conformed to the principles in the Declaration of Helsinki.

**Table 1.** Characteristics of the human participants. All values are presented as the mean ± SEM from *n* = 7-10 / group. The data for each of the variables was analyzed with two-way ANOVA, and the *P*-Values for the main effects and interactions are listed.

|  | Age<br>(yr) | Height<br>(cm) | Weight<br>(kg) | Body Fat<br>(%) | BMI | Femur<br>Length (cm) | Thigh Lean<br>Mass (kg) |
| --- | --- | --- | --- | --- | --- | --- | --- |
| Male Young | 27.1 ± 2.2 | 181.3 ± 2.7 | 84.1 ± 4.7 | 22.4 ± 1.3 | 25.4 ± 0.9 | 39.7 ± 1.5 | 8.8 ± 0.7 |
| Male Old | 74.3 ± 2.2 | 177.4 ± 2.3 | 83.4 ± 1.2 | 31.5 ± 1.8 | 26.6 ± 0.8 | 37.4 ± 0.6 | 6.3 ± 0.3 |
| Female Young | 23.0 ± 0.8 | 163.1 ± 3.4 | 63.1 ± 3.5 | 30.1 ± 2.6 | 23.7 ± 1.1 | 34.9 ± 0.5 | 5.4 ± 0.3 |
| Female Old | 69.0 ± 0.9 | 160.5 ± 2.1 | 65.0 ± 4.8 | 36.3 ± 2.6 | 25.1 ± 1.5 | 34.6 ± 0.6 | 4.5 ± 0.2 |
| Age Main Effect | *P* < 0.0001 | *P* = 0.2489 | *P* = 0.8733 | *P* = 0.0010 | *P* = 0.2168 | *P* = 0.1850 | *P* = 0.0007 |
| Sex Main Effect | *P* = 0.0146 | *P* < 0.0001 | *P* < 0.0001 | *P* = 0.0054 | *P* = 0.1309 | *P* = 0.0006 | *P* < 0.0001 |
| Interaction | *P* = 0.7369 | *P* = 0.8157 | *P* = 0.7450 | *P* = 0.5020 | *P* = 0.8935 | *P* = 0.3111 | *P* = 0.0834 |

### MRI of the knee extensor muscles

To quantify the volume and largest cross-sectional area (CSA) of the knee extensor muscles, participants laid supine in a 3.0 Tesla supraconducting magnet (Signa Premier, GE, Madison, WI, USA). The scanner operated with a 3D dual-echo, fat-water Dixon pulse sequence in the axial plane, yielding 7 mm thick serial cross-sections with the following parameters: TR = 8.28 ms, echo time (TE) = 2.97 ms, imaging frequency = 127.8 MHz, flip angle = 5°. Participants remained supine for at least 20 min prior to scanning to eliminate the effects of postural shifts in fluid volume on MRI measurements [24]. Axial slices from the T1-weighted water images were analyzed with Medical Image Processing, Analysis and Visualization (MIPAV, National Institutes of Health Center for Information Technology, version 11.2.0) beginning at the first slice where the patella was no longer visible and continuing through the slice corresponding to 80% of the total femur length. Femur length was measured as the distance between the first slice that includes the full tibial plateau and the last slice where the most proximal portion of the femoral head was still visible. For each slice, the knee extensor muscles (vastus medialis, vastus lateralis, vastus intermedius, and rectus femoris) were manually outlined if visible, and the corresponding volumes were quantified based on the CSA and slice thickness. The volumes for each of the four muscles were summed across all slices to determine muscle-specific volumes, and the total knee extensor muscle volume was determined as the combined volume of all four muscles. For ease of comparison to prior aging skeletal muscle literature, we also report the single 7 mm thick slice with the largest knee extensor CSA.

### Collection and processing of human skeletal muscle biopsies

A muscle biopsy from the vastus lateralis was obtained from each participant as described previously [25]. Briefly, participants were instructed to refrain from strenuous exercise of the lower limbs for 48 hr prior to the biopsy and arrive at the laboratory fasted and without consumption of caffeine for ≥8 hr. The biopsy site was cleaned with 70% ethanol, sterilized with 10% povidone-iodine, and anesthetized with 1% lidocaine HCl. A small incision, approximately 1 cm in length, was made over the distal third of the muscle belly, and the biopsy needle was inserted under local suction to obtain the tissue sample. A muscle bundle was aligned with the fibers oriented longitudinally on a small notecard presoaked in 0.1 M phosphate buffer (PB), and the sample was submerged in 0.1 M PB for 30 s to minimize potential dehydration of the bundle before undergoing tissue fixation and cryoprotection. To preserve the ultrastructural integrity of the myofibrils, we extensively fixed the sample in 4% paraformaldehyde and subsequently cryoprotected it with 15–45% sucrose as described previously [26]. Specifically, muscle bundles were fixed in 4% paraformaldehyde in 0.1 M PB for 24 hr (3 hr at 20°C on an orbital shaker at 50 RPM and 21 hr at 4°C on a nutating rocker), and then cyoprotected by incubating in 15% sucrose in 0.1 M PB for 6 hr (4°C) followed by 45% sucrose in 0.1 M PB for an additional 18 hr (4°C). The fixed and cryoprotected sample was then immersed in optimal cutting temperature (OCT) compound, frozen in liquid nitrogen-chilled isopentane, and stored at –80°C until sectioning.

### Animals

Experimental procedures were conducted on adult (4 months old, obtained from Jackson Laboratory) and aged (24 months old, obtained from the National Institute on Aging rodent colonies) male C57BL/6 mice. These animals were housed at 25 °C under a 12-hour light/dark cycle with free access to standard rodent chow and water, but no additional provisions for exercise enrichment such as running wheels or climbing devices. After shipment to University of Wisconsin-Madison, the mice were allowed a one-week acclimation period, and prior to immobilization, the mice were anesthetized with 2-5% isoflurane in oxygen. For tissue collection, animals were euthanized by cervical dislocation under anesthesia, followed by removal of the hindlimbs. All procedures were approved by the Institutional Animal Care and Use Committee (IACUC) of the University of Wisconsin-Madison (protocol #V005375).

### Hindlimb immobilization

Unilateral hindlimb immobilization was performed as previously described [27]. Briefly, mice were anesthetized with 2-5% isoflurane in oxygen, and the right hindlimb was immobilized with the knee extended and the ankle plantar-flexed for 10 days. The contralateral (left) hindlimb remained unrestrained and served as a control. Following the immobilization period, mice were anesthetized, and various tissues from the hindlimbs were collected as described below.

### Collection and processing of mouse skeletal muscles

Hindlimb collection and tissue processing were performed as previously described [26]. Briefly, mice were anesthetized with 2-5% isoflurane in oxygen, and then the skin on the hindlimbs was retracted, and the hindlimbs were severed at the hip joint. Next, for each hindlimb, the foot and proximal femur were secured to a circular aluminum wire mesh at 90° angles between the foot-tibia and tibia-femur joints using non-absorbable silk sutures, and then they were fixed in 20 mL of 4% paraformaldehyde in 0.1 M PB for 3 hr at room temperature on an orbital shaker at 50 RPM. Various muscles and the tibia from each hindlimb were then dissected, the length of the tibias and mass of the muscles were recorded, and then the soleus and flexor digitorum longus muscles were transferred to 1.5 mL tubes containing 1.0 mL of 4% paraformaldehyde in 0.1 M PB, and post-fixed for 21 hr at 4 °C on a nutating rocker. The muscles were then cryoprotected and frozen in OCT as described for the human samples.

### Immunohistochemistry

Mid-belly (mouse) and biopsy (human) muscle cross-sections (5 μm thick) were obtained from OCT-embedded tissue using a cryostat set to –30 °C and mounted on Superfrost Plus microscope slides (Fisher Scientific). Sections were immediately hydrated in diH2O for 15 min. While maintaining hydration, the area surrounding each section was gently dried with a Kimwipe, and a hydrophobic barrier was drawn around the tissue using a PAP pen (Aqua Hold 2, Scientific Device Laboratory #9804-02). Slides were then placed into a humidified chamber, where all subsequent washing and incubation steps were performed at room temperature on an orbital shaker at 50 RPM.

Sections were washed in phosphate-buffered saline (PBS) for 5 min and then incubated for 30 min in blocking solution (0.5% Triton X-100, 0.5% bovine serum albumin in PBS). For mouse samples, sections were incubated overnight with mouse IgG1 anti-SERCA1 (1:100, VE121G9, Santa Cruz #SC-58287) and rabbit anti-dystrophin (1:100, Thermo Fisher #PA1-21011) diluted in blocking solution. For human biopsy samples, sections were incubated overnight with either mouse IgG1 anti-SERCA1 or mouse IgG1 anti-SERCA2 (1:200, IID8, Santa Cruz #SC-53010) along with rabbit anti-dystrophin. After the overnight incubation, sections were washed three times for 10 min in PBS, followed by three additional washes for 30 min in PBS. The sections were then incubated overnight in blocking solution containing Alexa Fluor 594-conjugated goat anti-mouse IgG (Fcγ Subclass 1 specific; 1:2000, Jackson ImmunoResearch #115-585-205) and Alexa Fluor 488-conjugated goat anti-rabbit IgG (1:5000, Invitrogen #A11008). After incubation, the sections were washed three times for 10 min in PBS, followed by three additional washes for 30 min in PBS. The slides were then mounted with 4 μL of ProLong^TM^ Gold Antifade Mountant (Thermo Fisher #P36930) and covered with a No. 1 glass coverslip. Mounted sections were cured in the dark for at least 24 hr before imaging.

### Assessment of whole muscle size and muscle fiber size

Human and mouse cross-sections labeled for dystrophin and SERCA1/2 were imaged using a Keyence BZX700 automated inverted epifluorescence microscope with a 10× objective lens. For each fluorescent channel (FITC and TxRED), image fields were acquired as either 3×3 (mouse soleus samples) or 5×5 (mouse flexor digitorum longus and human samples) fields and stitched using Keyence Analyzer Software. Whole-muscle cross-sectional area (CSA) in mice and the mean CSA of the fibers in both human and mouse samples were quantified using our previously published CellProfiler pipeline [28].

### Fluorescence imaging of myofibrils and image deconvolution (FIM-ID)

Fluorescence imaging of myofibrils and image deconvolution (FIM-ID) were performed following our previously described method [26]. In brief, randomly selected regions of interest (ROIs) from cross-sections labeled for dystrophin and SERCA1/2 were imaged using a Leica THUNDER Imager Tissue 3D microscope with an HC PL APO 63×/1.40 oil immersion objective. Z-stacks were acquired across the full sample thickness and processed using Leica’s Small Volume Computational Clearing (SVCC) algorithm for deconvolution. The deconvolution images were then opened in ImageJ, and the Z-plane with the most in-focus image for SERCA1/2 was selected. The corresponding autofluorescence signal from the same plane was merged with the SERCA1/2 images using the Z-Project function in ImageJ, and the resulting merged image was saved as a single grayscale TIFF image. The final merged images were converted to 16-bit TIFF format, and the pixel density was standardized to 6144 × 6144 using the ImageJ resize function.

### Automated measurements of myofibril size and number with FIM-ID

For mouse samples, individual fibers were randomly selected from each FIM-ID image by a blinded investigator and manually traced in ImageJ to measure fiber CSA, as previously described [26]. Preliminary analyses included 15-30 fibers per sample (15 fibers per soleus samples and 30 fibers per flexor digitorum longus samples), with additional fibers randomly selected until the average CSA of the randomly selected fibers was within ± 10% of the mean fiber CSA obtained from the Keyence-based analyses of the whole muscle section. Single-fiber images of the signal for SERCA1 were then subjected to automated measurements of myofibril CSA with our “Myofibril CSA Analysis” pipeline in CellProfiler version 4.2.1 [26]. To estimate the number of myofibrils per fiber, each SERCA1 fiber image was processed with our “Intermyofibrillar Area” pipeline in CellProfiler, which quantifies the area in each fiber that is occupied by intermyofibrillar components (e.g., sarcoplasmic reticulum and mitochondria). The intermyofibrillar area was then subtracted from the total fiber CSA to yield the myofibrillar area, which was then divided by the mean myofibril CSA to determine the number of myofibrils per fiber. In some instances, the analyzed fibers were also grouped according to their oxidative capacity as previously described [26].

For human samples, the same general method was used with two modifications. First, analyses were performed on images labeled for SERCA1 or SERCA2. Second, preliminary analyses included 30 fibers (15 SERCA1 and 15 SERCA2), with additional fibers randomly selected until the average CSA was within ± 5% of the mean fiber CSA obtained from Keyence-based analyses of the whole biopsy section.

### Assessment of Tubular Aggregates

Randomly selected FIM-ID images of SERCA1 from young and old human vastus lateralis, mouse soleus, and mouse flexor digitorum longus muscles were assessed for the proportion of tubular aggregate-positive fibers by blinded investigators. Briefly, for each FIM-ID image, all SERCA1-positive fibers that were in focus and entirely encompassed within the image were counted. Then, the number of these fibers that contained tubular aggregates was determined. Three FIM-ID images per sample were analyzed, and the total number of tubular aggregate-positive and negative fibers from the three images was used to determine the proportion of tubular aggregate-positive fibers for each sample (8-29 fibers per sample for the vastus lateralis, 32-62 fibers per sample for the soleus, and 49-76 fibers per sample for the flexor digitorum longus).

### Statistical analysis

As previously described, samples that deviated more than three times from the mean within a given group were excluded as outliers [26]. As specified in the figure legends, statistical significance was assessed using unpaired t-tests, two-way ANOVA, mixed two-way ANOVA, or repeated-measures two-way ANOVA with Fisher’s LSD post hoc comparisons. Differences were considered significant at *P* < 0.05. All analyses were conducted using GraphPad Prism version 10.0.0 (Windows).

## Results

### Aging-induced changes in the macroscopic structure of human skeletal muscle

To identify aging-induced changes in the macroscopic structure of human skeletal muscle, MRI images of the thigh were collected from young and old participants (Table 1). The MRI images were then used to determine the volume of the knee extensor muscles (i.e., the vastus medialis, vastus lateralis, vastus intermedius, and rectus femoris) and the largest cross-sectional area (CSA) along the length of these muscles. As shown in Figure 1, the results indicated that aging led to a 34% reduction in the knee extensor muscle volume and a 32% reduction in the largest CSA of the muscles. Moreover, the outcomes revealed that the effects of aging were not significantly influenced by the sex of the participants (Supplemental Fig. 1).

**Figure 1.**
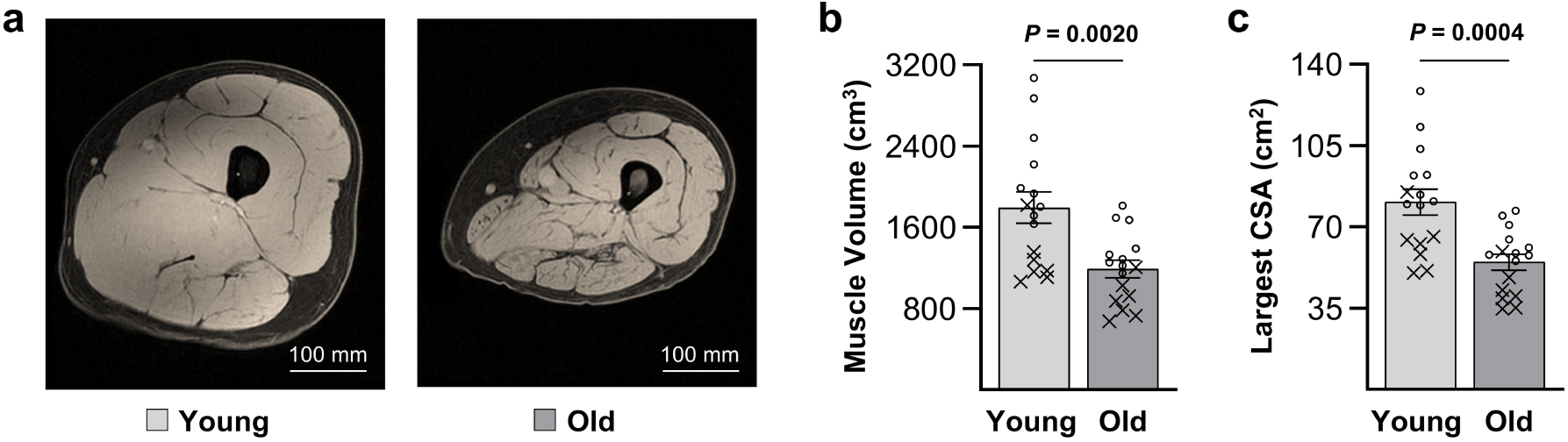
Aging-induced changes in the macroscopic structure of human skeletal muscle. (**a**) MRI images of the thigh were collected from young and old participants and used to determine (**b**) the volume of the knee extensor muscles (i.e., the vastus medialis, vastus lateralis, vastus intermedius, and rectus femoris) and (**c**) the largest cross-sectional area (CSA) along the length of the knee extensor muscles. The images in A are from male participants, scale bars = 100 mm. The data in B and C represent the combined results from male (○) and female (X) participants. Values are presented as group means ± SEM, *n* = 16 / group. The data were analyzed with unpaired t-tests, and the resulting *P*-values are displayed in the graphs.

### Aging-induced changes in the microscopic and ultrastructural properties of human skeletal muscle

Having established that aging led to a significant decrease in the volume and CSA of the knee extensor muscles, we next sought to identify the microscopic and ultrastructural adaptations that mediated these effects. To accomplish this, biopsies from the vastus lateralis of the participants were collected, and then cross-sections were subjected to immunohistochemistry for dystrophin to identify the periphery of the muscle fibers, and for SERCA1 or SERCA2 to identify the periphery of the myofibrils in Type II and I fibers, respectively (Fig. 2a-b) [26, 29]. With respect to microscopic adaptations, the outcomes revealed that aging led to radial atrophy of the muscle fibers (i.e., a decrease in fiber CSA) and that this effect was dependent on the fiber type. Specifically, aging led to a 23% decrease in the CSA of the SERCA1-positive fibers, but it did not significantly alter the CSA of the SERCA2-positive fibers (Fig. 2c).

**Figure 2.**
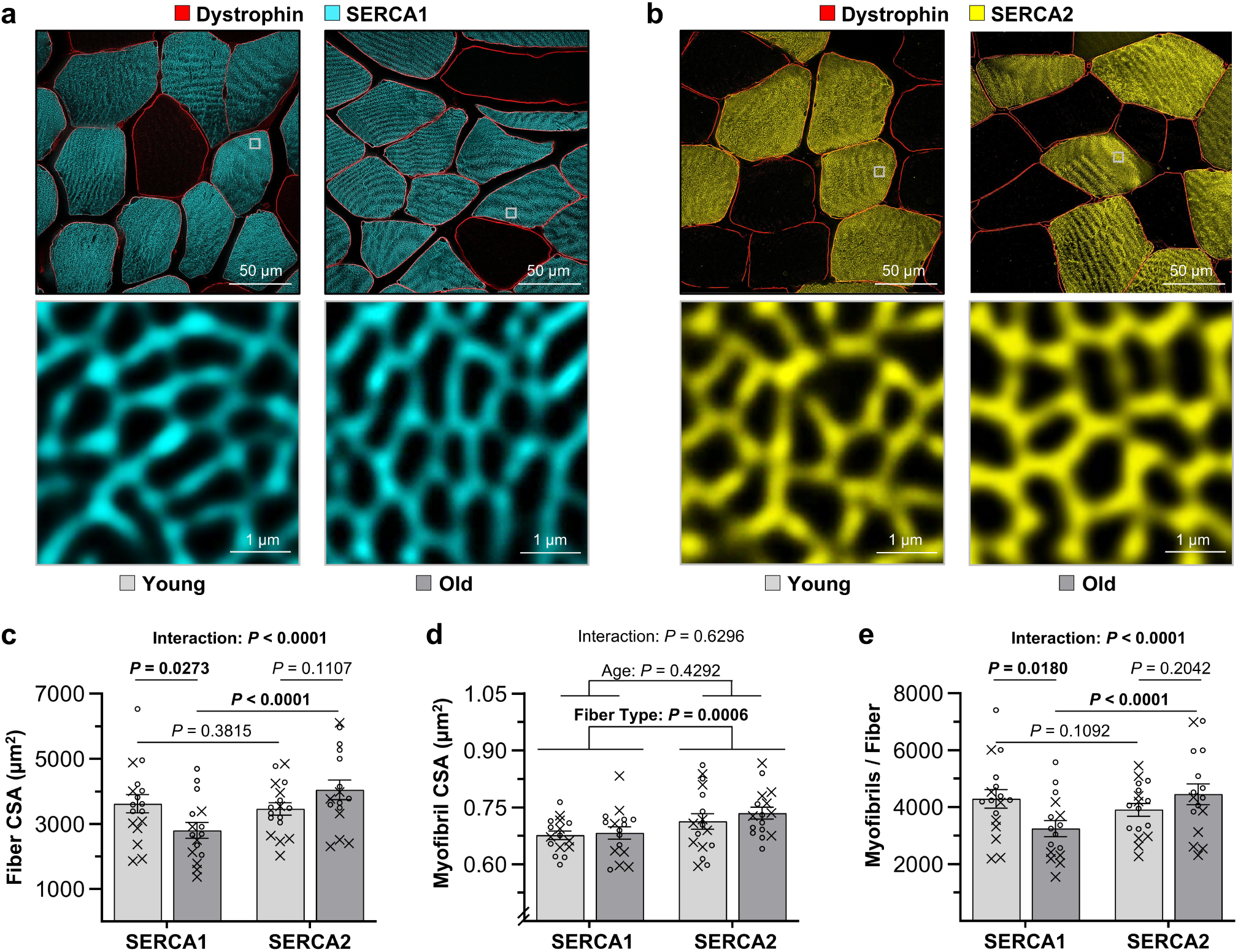
Aging-induced changes in the microscopic and ultrastructural properties of human skeletal muscle. Biopsies of the vastus lateralis of young and old participants were collected and then cross-sections were subjected to immunohistochemistry for dystrophin (to identify the periphery of muscle fibers) and (**a**) SERCA1 (to identify the periphery of the myofibrils in Type II fibers), or (**b**) SERCA2 (to identify the periphery of the myofibrils in Type I fibers). Scale bars = 50 μm in the low magnification images and 1 μm in the higher magnifications images of the boxed regions. The images were used to determine the average (**c**) cross-sectional area (CSA) of the fibers, (**d**) CSA of the myofibrils, and (**e**) number of myofibrils per fiber. The images in a-b are from male participants, whereas the values in c-e represent the combined results from males (○) and females (X). Values are presented as group means ± SEM, *n* = 16-17 muscles / group, 30-36 fibers / muscle, and an average of 957-1113 myofibrils per fiber. The data were analyzed with repeated measures (c, e) or mixed (d) two-way ANOVA, and the resulting *P*-values for the main effects, interactions, and pairwise comparisons are displayed in the graphs.

After determining that aging led to radial atrophy of the SERCA1-positive fibers, we sought to gain further insight into the ultrastructural adaptations that drove this effect. Specifically, we used our previously described FIM-ID pipeline to measure the average CSA of the myofibrils, as well as the average number of myofibrils per fiber [26]. As shown in Figure 2d, the outcomes revealed that aging did not significantly alter the CSA of the myofibrils in SERCA1- or SERCA2-positive fibers. However, aging did lead to a significant reduction in the number of myofibrils per fiber, and just like the changes in fiber CSA, this effect was specific to the SERCA1-positive fibers (Fig. 2e). As shown in Supplemental Figure 2, it was also determined that the aging-induced changes in the microscopic and ultrastructural properties of the SERCA1-positive fibers were not significantly influenced by the sex of the participants.

### The effects of aging and disuse on the mass of mouse skeletal muscles

Next, we wanted to examine whether the macro-to-ultrastructural changes observed with aging in human skeletal muscle were conserved in mice. However, the use of aged mice is expensive, and therefore, we designed our experiments so that we could maximize their use. For instance, the results in humans indicated that the effects of aging were not significantly influenced by sex; thus, we focused our studies on a single sex, and we chose males because previous studies have reported that skeletal muscles of aged male mice more closely recapitulate the aging phenotype of human skeletal muscle [30]. Furthermore, as illustrated in Figure 3a, we subjected the young (4-months of age) and old (24-months of age) mice to a model of unilateral hindlimb immobilization, which allowed us to assess both the effects of aging alone, as well as how aging influences the loss of muscle mass that occurs in response to disuse. Specifically, at 10 days after the onset of the immobilization, body mass and the length of the tibia in each hindlimb were recorded (Fig. 3b-c), and then the tibialis anterior (TA), extensor digitorum longus (EDL), plantaris (PLT), soleus (SOL), gastrocnemius (GAST), and flexor digitorum longus (FDL) muscles from each hindlimb were weighed and saved for additional analyses (Fig. 3d-e).

**Figure 3.**
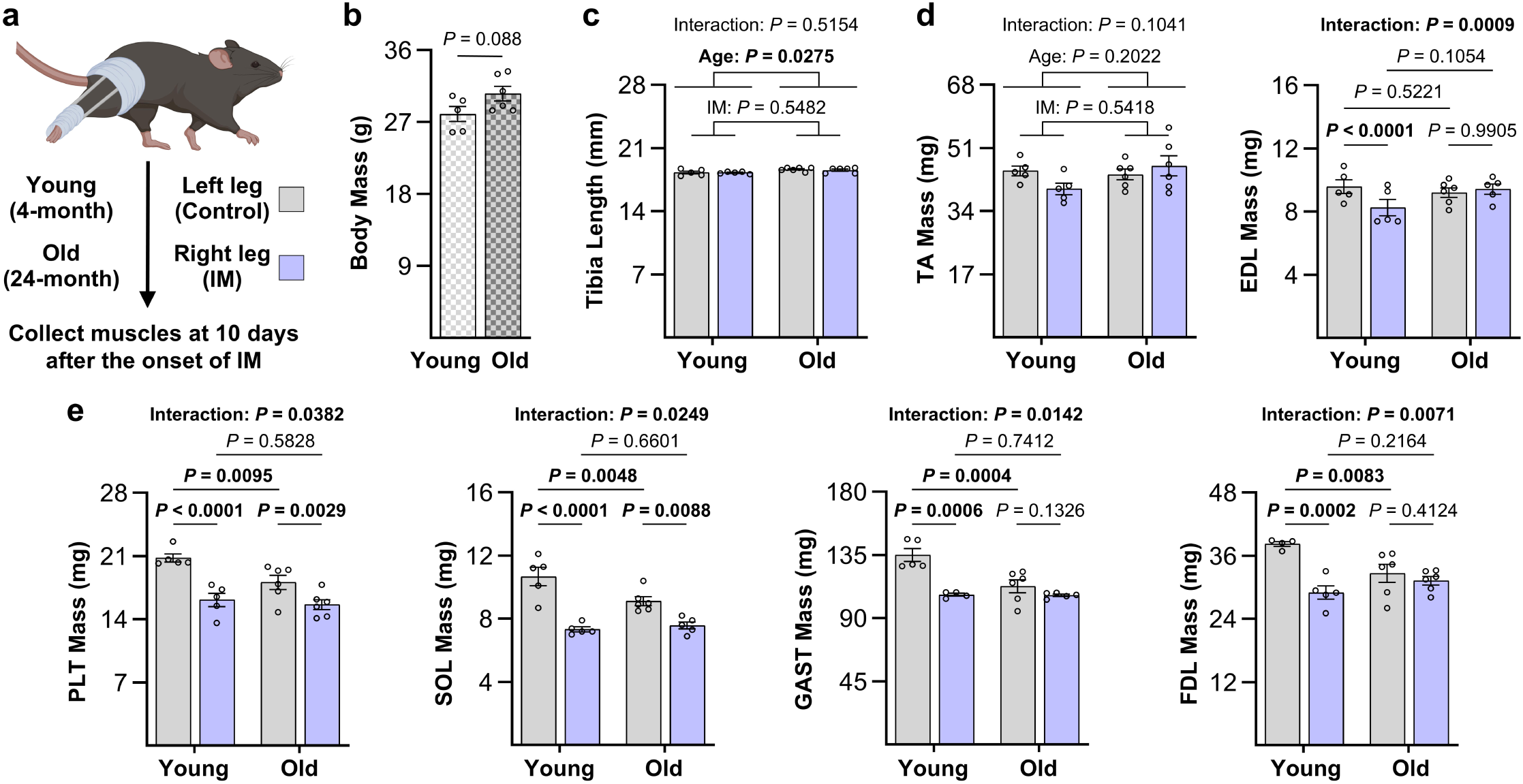
The effects of aging and disuse on the mass of various mouse skeletal muscles. (**a**) Young and old male C57BL6 mice were subjected to disuse via unilateral immobilization (IM) of the right hindlimb, whereas the left limb served as a control (created with BioRender.com). After 10 days, measurements were obtained for body mass (**b**), tibia length (**c**), and the mass of individual hindlimb muscles including (**d**) extensors such as the tibialis anterior (TA), and extensor digitorum longus (EDL), and (**e**) flexors such as the plantaris (PLT), soleus (SOL), gastrocnemius (GAST), and flexor digitorum longus (FDL). Values are presented as group means ± SEM, *n* = 4-6 / group. The data were analyzed with an unpaired t-test (b), repeated measures two-way ANOVA (c, d), or mixed two-way ANOVA (d, e), and the resulting *P*-values for the main effects, interactions, and pairwise comparisons are displayed in the graphs.

To assess how immobilization and aging impacted muscle mass, the data were first subjected to a three-way ANOVA, and the outcomes revealed that there was a significant interaction between the effects of age and immobilization, and that the interaction was dependent on the muscle being examined (*P* = 0.0003). Accordingly, we performed two-way ANOVAs that assessed the effects of age and immobilization in each muscle, and the outcomes revealed that a significant interaction between these variables was present in all muscles except for the TA (Fig. 3d-e). Further analysis revealed that the interaction in these muscles could be attributed to a blunted immobilization-induced decrease in mass in the aged animals. Subsequent pair-wise analyses also revealed that all of the flexor muscles (i.e., the PLT, SOL, GAST, and FDL) exhibited a significant aging-induced loss of mass, but this effect was not present in either of the extensor muscles (i.e., the TA and EDL).

### Aging-induced changes in the macro-to-ultrastructural properties of mouse skeletal muscle

Having established that various muscles of mice exhibit an aging-induced loss of mass, we next set out to determine whether this effect was mediated by the same macro-to-ultrastructural changes that we observed in humans. Specifically, as shown in Figure 4, mid-belly cross-sections of the mixed slow/fast-twitch SOL (i.e., Type I and IIa fibers [31]), as well as the highly fast-twitch FDL (i.e., Type IIa, IIx, and IIb fibers [28]), were subjected to immunohistochemistry for dystrophin to identify the periphery of the muscle fibers, and SERCA1 to identify the periphery of the myofibrils in the Type II fibers (i.e., the fiber type that showed aging-induced radial atrophy in the human samples). Interestingly, during our analysis of the FDL muscles, we noticed non-myofibrillar anomalies in the signal for SERCA1, with both the number and size of these anomalies being greater in the FDL muscles from the aged mice (Fig. 4b). We suspected that these anomalies represented tubular aggregates which were originally described by Engel et al. (1970) as an abnormal accumulation of the sarcoplasmic reticulum [32]. As shown in Supplemental Figure 3, the presence of these anomalies also aligned with previous studies, which have shown that tubular aggregates are rare in healthy skeletal muscles of humans, but are prevalent in the fast-twitch fibers of mouse skeletal muscles, where aging leads to an increase in both their size and number [32, 33]. Importantly, since the presence of these anomalies would confound our assessment of myofibril CSA and myofibril number, all analyses for the FDL muscles were performed on fibers that did not contain the anomalies, yet were matched with the whole population in terms of their average CSA (see methods for details).

**Figure 4.**
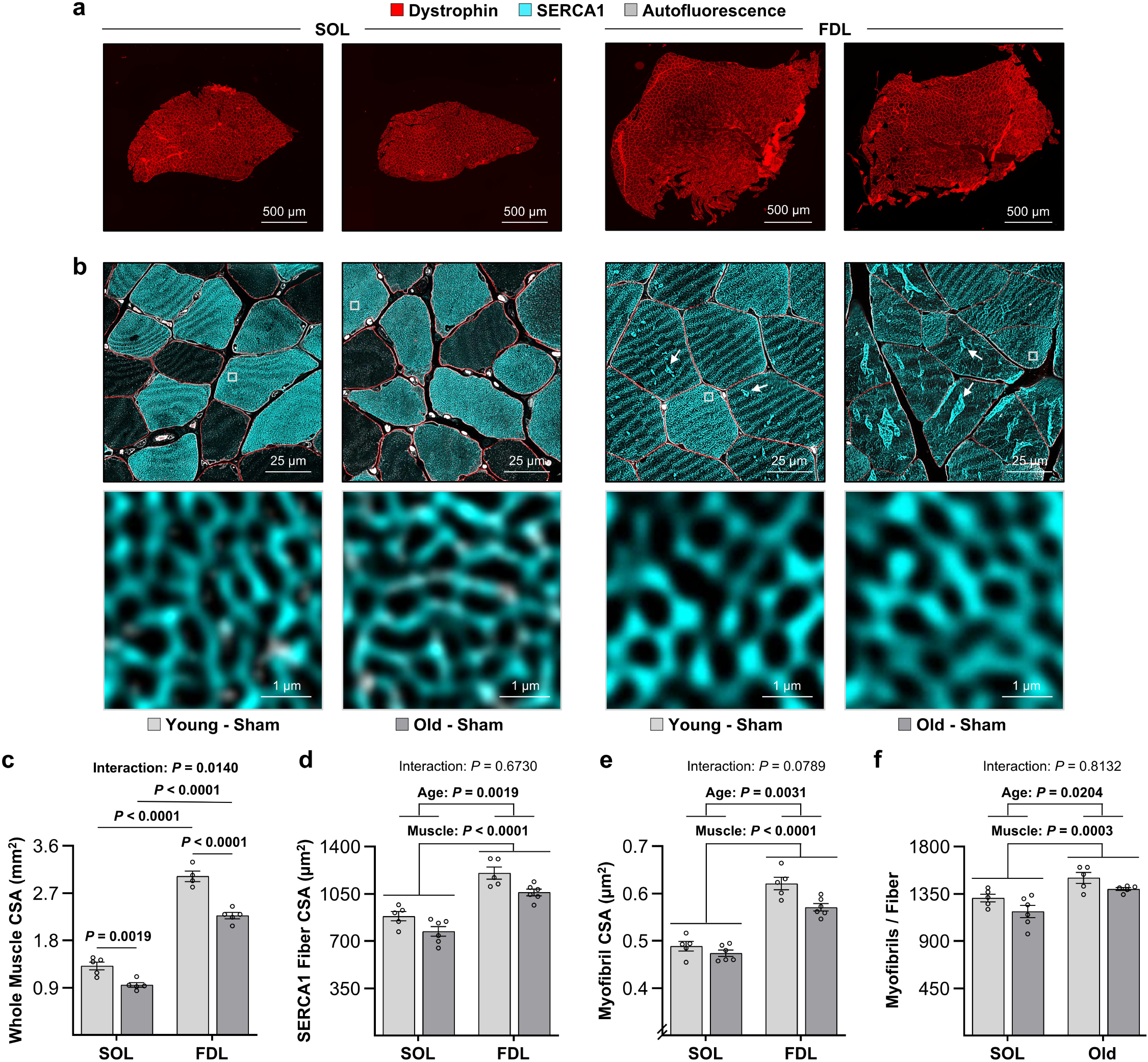
Aging-induced changes in the macro-to-ultrastructural properties of mouse skeletal muscle. Mid-belly cross-sections were obtained from the SOL and FDL muscles of the control limb of the mice described in Figure 3. The sections were subjected to immunohistochemistry for dystrophin (**a**), or dystrophin and SERCA1 (**b**), and imaged at various levels of magnification (scale bar a = 500 μm, b = 25 μm in the low magnification images and 1 μm in the higher magnifications images of the boxed regions). The resulting images were merged with the signal from autofluorescence and used to determine the (**c**) whole muscle cross-sectional area (CSA), (**d**) the average CSA of the SERCA1 positive fibers, as well as the (**e**) average CSA of the myofibrils, and (**f**) number of myofibrils per fiber in the SERCA1 positive fibers. The data were grouped by the specific muscle and age, and presented as group means ± SEM, *n* = 4-6 muscles / group, 15-34 fibers / muscle, and an average of 385-411 myofibrils per fiber. The data were analyzed with two-way ANOVA, and the resulting *P*-values for the main effects, interactions, and pairwise comparisons are displayed in the graphs. Arrows in b point to examples of the putative tubular aggregates.

As reported in Figure 4, the outcomes of our analyses revealed that the aging-induced loss of muscle mass in both the SOL and FDL was accompanied by a reduction in the whole muscle CSA. Similar to our observations in humans, we also found that aging led to radial atrophy of the SERCA1-positive fibers, and that this effect could be attributed, at least in part, to a reduction in the number of myofibrils per fiber. However, unlike in humans, aging also led to a small but significant decrease in the CSA of the myofibrils. Moreover, two-way ANOVA revealed that there was a trend for this to occur in a muscle-specific manner (3% vs. 8% in the SOL vs. FDL, respectively, *P* = 0.0789). Thus, while an aging-induced decrease in the number of myofibrils per fiber appears to be a conserved phenomenon, an aging-induced decrease in myofibril CSA might be an event that occurs specifically in mice and/or in a muscle-/fiber type-specific manner.

### The effect of aging on disuse-induced changes in the macro-to-ultrastructural properties of mouse skeletal muscle

After characterizing the macro-to-ultrastructural changes that are associated with the aging-induced loss of muscle mass, we then examined whether similar changes occur during the disuse-induced loss of muscle mass and whether these effects are influenced by aging. Specifically, mid-belly cross-sections were obtained from the SOL muscles of the control and immobilized limbs of both the young and old mice. The cross-sections were then subjected to immunohistochemistry for dystrophin to identify the periphery of the muscle fibers, and for SERCA1 to identify the periphery of the myofibrils (Fig. 5a-b). In line with the changes that were observed for muscle mass (Fig. 3e), the outcomes revealed that immobilization led to a significant decrease in the whole muscle CSA of the SOL in young mice, and that the magnitude of this effect was significantly reduced in the SOL muscles from the old mice (Fig. 5c). It was also determined that immobilization led to radial atrophy of the SERCA1-positive fibers and, just like the changes in whole muscle CSA, the magnitude of this effect was dependent on the age of the mice (Fig. 5d). Moreover, the radial atrophy of the SERCA1-positive fibers could be attributed to an immobilization-induced decrease in the CSA of the individual myofibrils, as well as to a decrease in the number of myofibrils per fiber (Fig. 5e-f). It also bears mentioning that nearly identical outcomes were observed when the same types of analyses were performed on FDL muscles (Supplemental Fig. 4).

**Figure 5.**
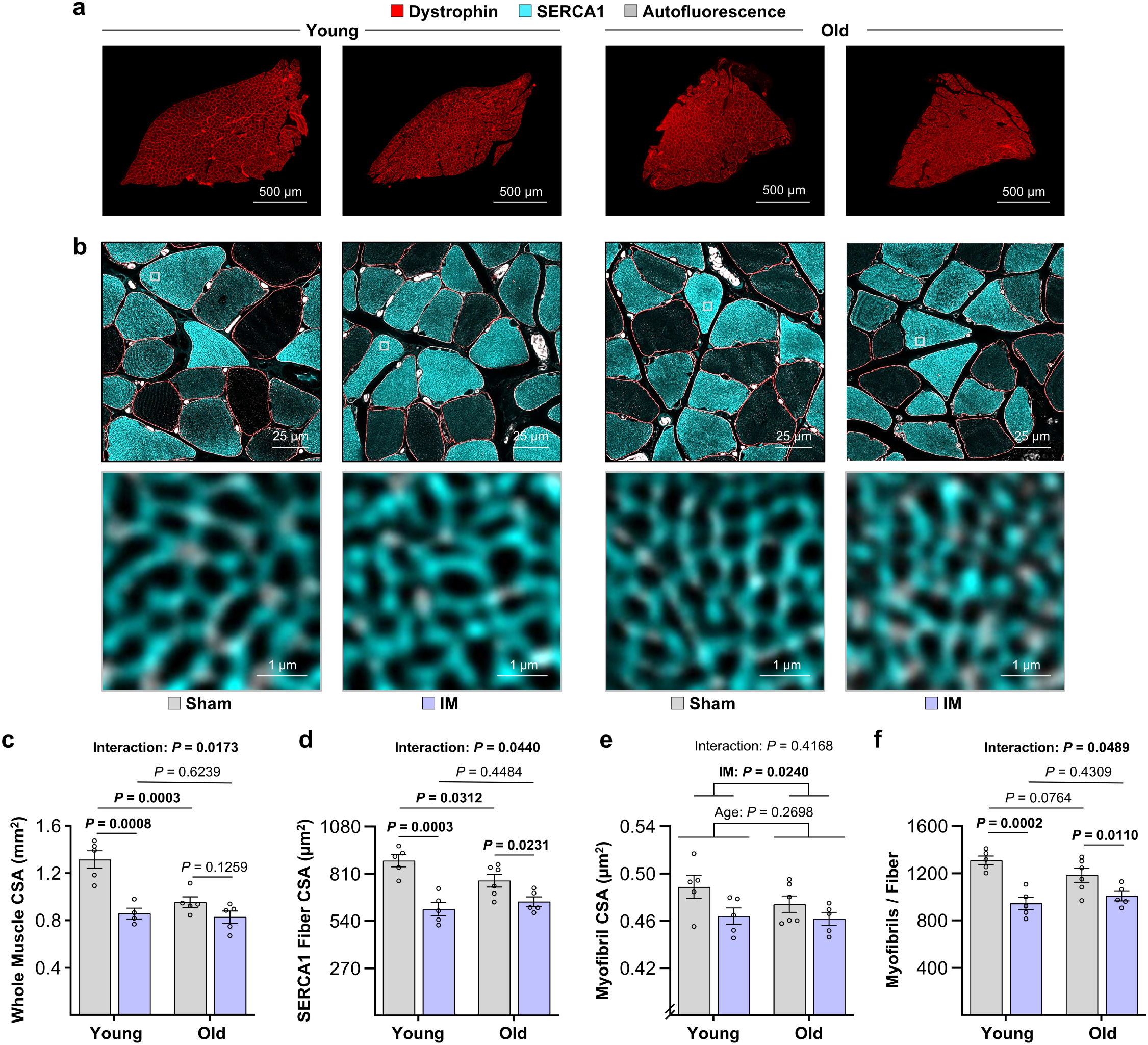
The effect of aging on disuse-induced changes in the macro-to-ultrastructural properties of the soleus muscle in mice. Mid-belly cross-sections were obtained from the SOL muscles of the mice described in Figure 3. The sections were subjected to immunohistochemistry for dystrophin (**a**), or dystrophin and SERCA1 (**b**), and imaged at various levels of magnification (scale bar a = 500 μm, b = 25 μm in the low magnification images and 1 μm in the higher magnifications images of the boxed regions). The resulting images were merged with the signal from autofluorescence and used to determine the (**c**) whole muscle cross-sectional area (CSA), (**d**) the average CSA of the SERCA1 positive fibers, as well as the (**e**) average CSA of the myofibrils, and (**f**) number of myofibrils per fiber in the SERCA1 positive fibers. The data were grouped by the specific muscle and age, and presented as group means ± SEM, *n* = 4-6 muscles / group, 15-17 fibers / muscle, and an average of 369-399 myofibrils per fiber. The data were analyzed with mixed two-way ANOVA, and the resulting *P*-values for the main effects, interactions, and pairwise comparisons are displayed in the graphs.

## Discussion

For nearly 50 years, studies have consistently concluded that the radial atrophy of muscle fibers is a major contributor to the loss of muscle mass that occurs during aging and disuse [14, 15, 20, 21]. However, despite the widespread recognition of this phenomenon, the ultrastructural adaptations that drive the radial atrophy remain poorly defined. One reason for this gap is that myofibrils are the dominant ultrastructural component of the muscle fibers, and rigorous quantification of their properties requires the tracing of thousands of myofibrils per sample [26]. This challenge is further compounded by the fact that myofibrils are usually imaged with electron microscopy, a technique that generally does not provide the level of contrast needed for automated myofibril tracing [26]. As such, assessments of myofibril properties have traditionally relied on manual myofibril tracing [26], and this means that even modestly powered studies could require hundreds to thousands of hours of labor. Thus, to address this limitation, we recently developed FIM-ID, a light microscopy-based approach that generates high-contrast images that are amenable to automated myofibril quantification [26].

With the advent of FIM-ID, our group was uniquely equipped with the ability to assess the macro-to-ultrastructural adaptations that mediate the aging- and disuse-induced loss of skeletal muscle mass. In humans, the primary outcomes from our analyses revealed that aging led to a reduction in knee extensor muscle volume and cross-sectional area that was accompanied by radial atrophy of SERCA1-positive fibers. Importantly, our ultrastructural analyses revealed that the radial atrophy of the fibers was driven by a decrease in the number of myofibrils per fiber, with no change in the CSA of the individual myofibrils. This is important because it suggests that the aging-induced radial atrophy of the muscle fibers is mediated by a defect in myofibril turnover, rather than a more generalized and uniform degradation of the existing contractile machinery.

Parallel experiments in mice largely recapitulated the findings in humans, with coordinated macro-to-ultrastructural declines occurring in multiple muscles during aging. However, unique to mice, the CSA of the myofibrils also showed signs of a decrease, particularly in the highly fast-twitch FDL muscle. The reason for this discrepancy is not known, but we suspect that it might, at least in part, be due to differences in the fiber-type composition of human and mouse skeletal muscles. Specifically, nearly 60% of the fibers in the FDL are glycolytic/Type IIb (an isoform that is not present in human skeletal muscle), and we determined that the CSA of the myofibrils in oxidative SERCA1-positive fibers of the FDL was not significantly altered by aging, whereas the CSA of the myofibrils in the glycolytic SERCA1-positive fibers experienced a clear reduction (Supplemental Figure 5) [28, 34]. These results are consistent with the notion that the highly glycolytic/Type IIb fibers exhibit a distinct vulnerability to aging. In further support of this, we also determined that tubular aggregates were highly prevalent in the FDL muscles of the old mice, but were rarely observed in the mixed slow/fast twitch SOL of the mice or the vastus lateralis of the humans. Indeed, previous studies have reported that tubular aggregates are found exclusively within Type IIb fibers of aged mice [33]. Thus, while our collective findings suggest that the overall trajectory of macro-to-ultrastructural adaptations that occur with aging are broadly conserved, there are also unique aging-induced adaptations that occur in muscles with highly glycolytic/Type IIb fibers.

The analyses performed in this study also provide insight into the mechanisms that mediate disuse-induced muscle atrophy and how these processes are altered with advancing age. Specifically, immobilization resulted in radial atrophy of the muscle fibers, and this effect was mediated by a reduction in both the CSA of the myofibrils and the number of myofibrils. These findings align with prior reports demonstrating rapid contractile protein loss during periods of inactivity [35, 36] and extend the literature by revealing that the structural basis for this loss involves remodeling at multiple ultrastructural levels. Notably, our analyses also revealed that the magnitude of disuse-induced changes was markedly blunted in aged muscle. However, this was not entirely unexpected, as previous studies have reported similar results [37, 38] and might simply reflect the fact that aged mice exhibit a substantial decline in daily physical activity [39]. Put differently, we suspect that many of the muscles in the aged animals were conditioned with chronic inactivity/disuse and were therefore less responsive to the additional atrophic effects of immobilization.

Although the comparative nature of our study is a major strength, there are also several limitations that need to be acknowledged. For instance, the analyses in humans were restricted to the vastus lateralis and therefore would not capture muscle-specific differences that likely exist across the body [40]. Second, the cross-sectional nature of our studies precludes definitive conclusions about causality or the temporal sequence of the structural adaptations. Third, while the immobilization model enabled mechanistic insights into disuse-induced atrophy, differences in fiber-type composition between species (particularly the presence of highly glycolytic/Type IIb fibers in mice) limit the extrapolation of certain findings to humans. Finally, although our use of FIM-ID provided robust ultrastructural measurements, it did not provide insight into the molecular mechanisms (e.g., signaling pathways, protein turnover rates, etc.) that drove the observed changes. Thus, additional studies will be needed to define these important mechanisms.

## Conclusion

This study provides a comprehensive macro-to-ultrastructural framework for understanding how aging and disuse lead to the loss of skeletal muscle mass. The results reveal that a reduction in myofibril number represents a central and conserved mechanism underlying radial atrophy, while changes in myofibril size contribute in a context- and muscle-/fiber type-specific manner. These findings not only clarify long-standing questions regarding the structural basis of muscle atrophy but also highlight potential targets for therapeutic interventions aimed at preserving muscle mass and function across the lifespan. Future work should focus on identifying the molecular pathways that regulate myofibril turnover and determining how these processes can be modulated to enhance muscle resilience in aging and disease.

## Acknowledgements

This work was supported by the National Institute on Aging grant AG048262 to CWS and the National Institute of Arthritis and Musculoskeletal and Skin Diseases grants AR08281 and AR057347 to TAH. We thank Kam Kroner and Lanny Klamecki for assisting with a portion of the MRI analysis.

## Ethics Statement

All human and animal studies were approved by the appropriate ethics committee and conformed to the principles in the Declaration of Helsinki.

## Data Availability Statement

All data are available upon reasonable request from the corresponding author.

---

**Supplemental Figure 1 Aging-induced changes in the macroscopic structure of human skeletal muscle are not influenced by sex** (**a**) MRI images of the thigh were collected from young and old participants and used to determine (**b**) the volume of the knee extensor muscles (i.e., the vastus medialis, vastus lateralis, vastus intermedius, and rectus femoris) and (**c**) the largest cross-sectional area (CSA) along the length of the knee extensor muscles. The images in A are from female participants, scale bars = 100 mm. The data in b and c were grouped by the age and sex of the participants and presented as group means ± SEM, with *n* = 7-9 / group. The data were analyzed with two-way ANOVA, and the resulting *P*-values for the main effects and interactions are displayed in the graphs.

**Supplemental Figure 2 Aging-induced changes in the microscopic and ultrastructural properties of SERCA1-positive fibers are not influenced by sex** (**a**-**b**) Biopsies of the vastus lateralis of young and old participants were collected, and then cross-sections were subjected to immunohistochemistry for dystrophin (to identify the periphery of muscle fibers) and SERCA1 (to identify the periphery of the myofibrils). Scale bars = 50 μm in the low magnification images and 1 μm in the higher magnifications images of the boxed regions. The images were used to determine the average (**c**) cross-sectional area (CSA) of the SERCA1 positive fibers, (**d**) CSA of the myofibrils, and (**e**) number of myofibrils per fiber. The data were grouped by the age and sex of the participants and presented as group means ± SEM, *n* = 6-9 muscles / group, 15-18 fibers / muscle, and an average of 957-1113 myofibrils per fiber. The data were analyzed with two-way ANOVA, and the resulting *P*-values for the main effects and interaction are displayed in the graphs. The images in a-b are from female participants.

**Supplemental Figure 3 Aging leads to an increase in tubular aggregates in the FDL muscle of mice** (**a**) Representative image of the FDL from an old mouse that was subjected to immunohistochemistry for dystrophin (red) and SERCA1 (cyan), scale bar = 15 µm. (**b**) An autofluorescent image of the same region shown in a reveals that the areas with intense SERCA1 staining (i.e., the tubular aggregates) exhibit a decrease in autofluorescence. (**c**) The images of SERCA1 positive fibers in the vastus lateralis (VL) of young (Y) and old (O) humans (Fig. 2), as well as the images of SERCA1 positive fibers from the control soleus (SOL) and control flexor digitorum longus (FDL) of Y and O mice (Fig. 4), were scored for the presence of tubular aggregates. Values are presented as group means ± SEM, *n* = 4-6 / group. The data were analyzed with two-way ANOVA, and the resulting *P*-values for the interaction and pairwise comparisons are displayed in the graphs.

**Supplemental Figure 4 The effect of aging on disuse-induced changes in the macro-to-ultrastructural properties of the FDL muscle in mice** Mid-belly cross-sections were obtained from the FDL muscles of the mice described in Figure 3, processed as in Figure 5, and used to determine the (**a**) whole muscle cross-sectional area (CSA), (**b**) the average CSA of the SERCA1 positive fibers, as well as the (**c**) average CSA of the myofibrils, and (**d**) number of myofibrils per fiber in the SERCA1 positive fibers. The data were grouped by the specific muscle and age, and presented as group means ± SEM, *n* = 4-6 muscles / group, 30-34 fibers / muscle, and an average of 368-411 myofibrils per fiber. The data were analyzed with mixed two-way ANOVA, and the resulting *P-*values for the main effects, interactions, and pairwise comparisons are displayed in the graphs.

**Supplemental Figure 5 The effect of aging on the CSA of myofibrils in the oxidative versus glycolytic SERCA1-positive fibers of the mouse FDL** Mid-belly cross-sections of the FDL muscles were subjected to immunohistochemistry for dystrophin and SERCA1 and subsequently imaged as described in Figure 4. The autofluorescence signal was used to classify the SERCA1-positive fibers as being oxidative (OX) or glycolytic (GLY), and the mean CSA of the myofibrils for each fiber type was then determined. The resulting data were grouped by the specific fiber type and age, and presented as group means ± SEM, *n* = 4-6 muscles / group, 30-34 fibers / muscle, and an average of 343-401 myofibrils per fiber. The data were analyzed with unpaired t-tests, and the resulting *P*-values are displayed in the graphs.

