## Supplemental Figures 1-5 for "Macroscopic to Ultrastructural Analyses Identify the Loss of Myofibrils as the Primary Mediator of Muscle Fiber Atrophy in Aging and Disuse"

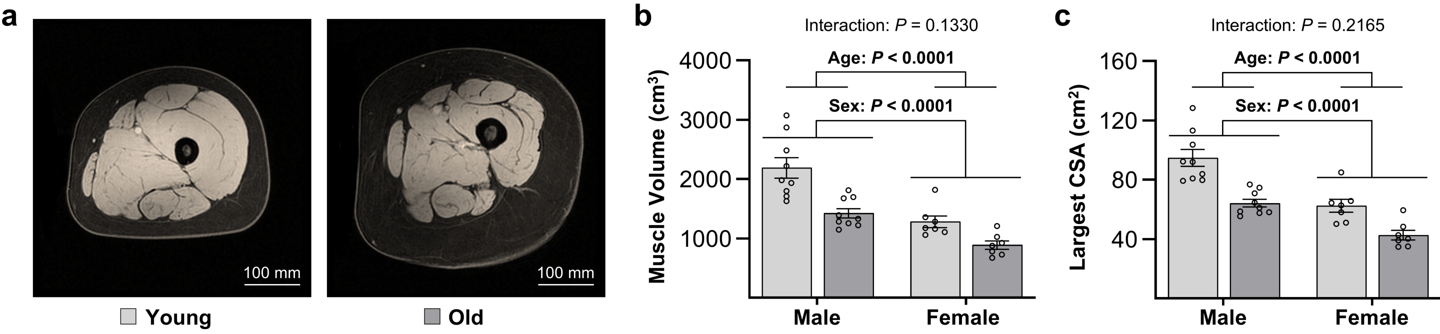


### Supplemental Figure 1

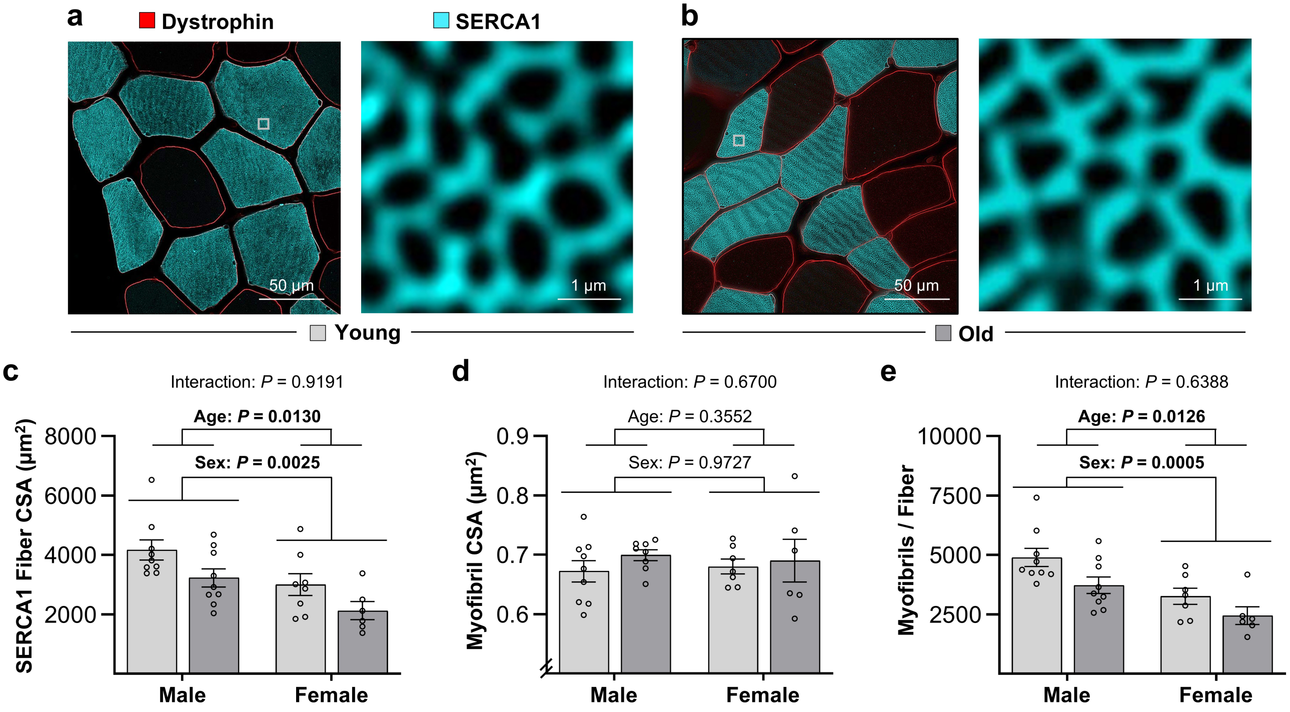


**Supplemental Figure 2**

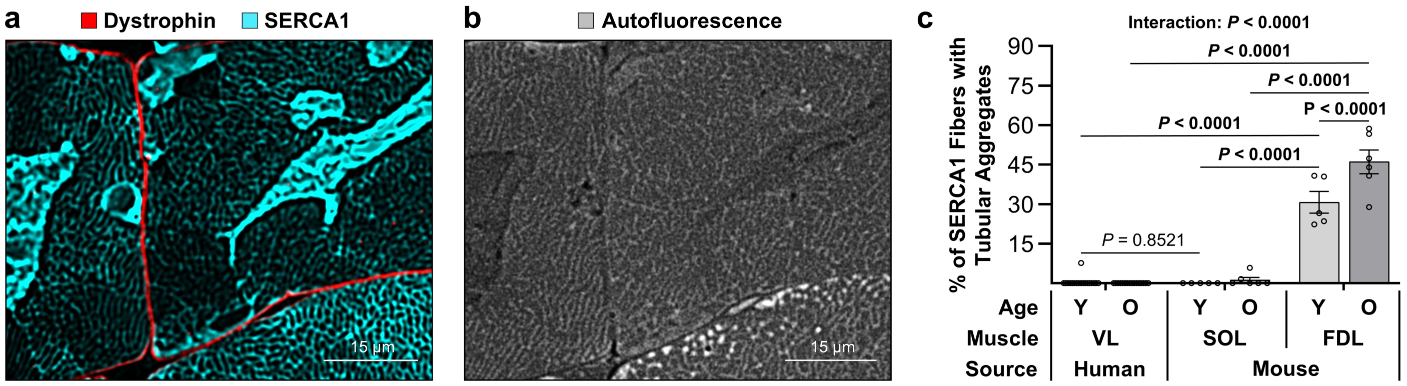


### Supplemental Figure 3

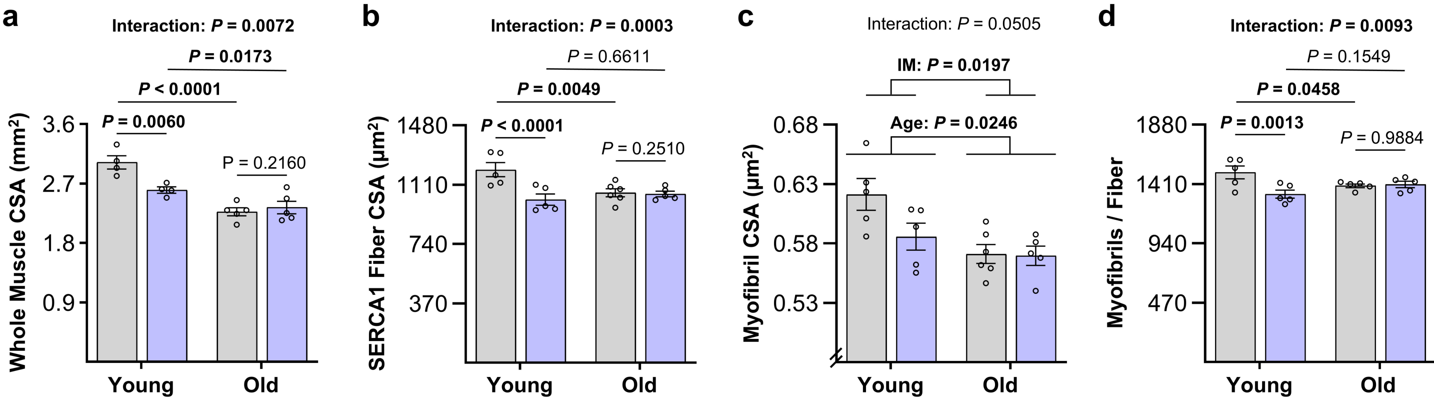


### Supplemental Figure 4

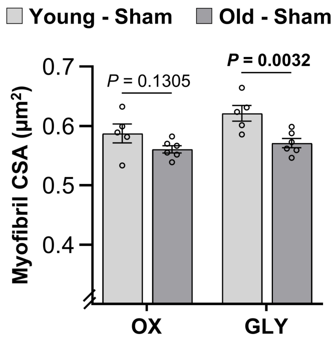


### Supplemental Figure 5
